# Reproducible Sex Differences in Personalized Functional Network Topography in Youth

**DOI:** 10.1101/2024.09.26.615061

**Authors:** Arielle S. Keller, Kevin Y. Sun, Ashley Francisco, Heather Robinson, Emily Beydler, Dani S. Bassett, Matthew Cieslak, Zaixu Cui, Christos Davatzikos, Yong Fan, Margaret Gardner, Rachel Kishton, Sara L. Kornfield, Bart Larsen, Hongming Li, Isabella Linder, Adam Pines, Laura Pritschet, Armin Raznahan, David R. Roalf, Jakob Seidlitz, Golia Shafiei, Russell T. Shinohara, Daniel H. Wolf, Aaron Alexander-Bloch, Theodore D. Satterthwaite, Sheila Shanmugan

## Abstract

**Background:** A key step towards understanding psychiatric disorders that disproportionately impact female mental health is delineating the emergence of sex-specific patterns of brain organization at the critical transition from childhood to adolescence. Prior work suggests that individual differences in the spatial organization of functional brain networks across the cortex are associated with psychopathology and differ systematically by sex.

**Aims:** We aimed to evaluate the impact of sex on the spatial organization of person-specific functional brain networks.

**Method:** We leveraged person-specific atlases of functional brain networks defined using non-negative matrix factorization in a sample of *n* = 6437 youths from the Adolescent Brain Cognitive Development Study. Across independent discovery and replication samples, we used generalized additive models to uncover associations between sex and the spatial layout (“topography”) of personalized functional networks (PFNs). Next, we trained support vector machines to classify participants’ sex from multivariate patterns of PFN topography. Finally, we leveraged transcriptomic data from the Allen Human Brain Atlas to evaluate spatial correlations between sex differences in PFN topography and gene expression.

**Results:** Sex differences in PFN topography were greatest in association networks including the fronto-parietal, ventral attention, and default mode networks. Machine learning models trained on participants’ PFNs were able to classify participant sex with high accuracy. Brain regions with the greatest sex differences in PFN topography were enriched in expression of X-linked genes as well as genes expressed in astrocytes and excitatory neurons.

**Conclusions:** Sex differences in PFN topography are robust, replicate across large-scale samples of youth, and are associated with expression patterns of X-linked genes. These results suggest a potential contributor to the female-biased risk in depressive and anxiety disorders that emerge at the transition from childhood to adolescence.

## Introduction

Many psychiatric disorders show sex differences in prevalence, presentation and trajectory. For example, the lifetime prevalence of internalizing disorders such as depression and anxiety is nearly twice as high in females^1^, and developmental disorders such as attention-deficit hyperactivity disorder often present differently in males and females leading to disparities in diagnosis and treatment. These sex differences tend to emerge during the transition from childhood to adolescence, a time when functional brain networks implicated in these disorders are refined^2,3^. Understanding and treating mental health conditions that are more prevalent in and differentially impact females requires a clear understanding of sex differences in neurodevelopment.

Prior neuroimaging studies have revealed significant sex differences in functional networks supporting cognitive and emotional processes, including the fronto-parietal^4,5^ and default mode^6^ networks. Dysfunction within these networks has been linked with psychiatric disorders, including anxiety and depression^7–11^. Critically, these functional networks are highly person-specific in their spatial organization across the cortex (“functional topography”). Substantial individual differences in the size, shape, and spatial location of brain regions comprising these networks emerge gradually during neurodevelopment with evidence of sex-specific patterning^3,12^. Innovations in precision brain mapping approaches have begun to chart the person-specific functional topography of personalized functional brain networks (PFNs)^13–15^ and have uncovered novel associations with internalizing psychopathology^11,16,17^ and cognition^3,18^. In a recent study of individuals across a broad age range (*n=*693, 8-22 years old)^12^, we presented the first report of sex differences in PFN functional topography. Given the ongoing reproducibility crisis in psychology and neuroscience^19^, it is important to determine whether such effects generalize across diverse samples. Moreover, it remains unclear whether these sex differences are consistently observed at the critical transition from childhood to adolescence when many psychiatric disorders first emerge. Here, we examine sex differences in PFN topography in youth using non-linear modeling, machine learning, and imaging transcriptomics in data from the Adolescent Brain Cognitive Development (ABCD) Study^®20^ (*n*=6,437, ages 9-10). We hypothesized that sex differences would be greatest in association networks and these sex differences would spatially correlate with X-linked gene expression.

## Method

### Participants

Participants from the ABCD Study^®20^ baseline assessment were drawn from the ABCD BIDS Community Collection (ABCC, ABCD-3165^21^). These data were collected across 22 sites in the United States, with Institutional Review Board (IRB) approval from the University of California, San Diego, as well as each of the respective study sites. Written informed consent (parents or guardians) and assent (children) were obtained. Criteria for participation in the ABCD Study^®^ are described in detail in previous work^22^. From the full baseline sample (*n*=11,878, 9-10 years old), we excluded participants with incomplete data or excessive head motion during fMRI scanning (**Figure S1**). Analyses were conducted in matched discovery (*n*=3240, 50.46% female) and replication (*n*=3197, 49.13% female) samples drawn from the ABCD Reproducible Matched Samples (ARMS^21,23^). We excluded siblings separately in the discovery and replication samples to avoid leakage across subsamples during model cross-validation (**Figure S1**). Importantly for the present study, we note that participant “sex” was assessed using a binary caregiver-reported question regarding the assignment of sex at birth on the original birth certificate. Hereafter, we use the term “sex” to refer to sex assigned at birth, the term “female” to refer to individuals assigned female at birth, and the term “male” to refer to individuals assigned male at birth.

### Definition of personalized functional networks (PFNs)

Details of the neuroimaging acquisition for the ABCD Study^®24^ and our fMRI preprocessing steps in this sample have been described previously^18,25^ (**Supplemental Information**).

Functional brain regions comprising large-scale networks have been shown to vary substantially in their size, shape, and spatial location across individuals^13,14^. We therefore employed a precision brain mapping approach as in previous work^3,12,16,18,25^ that leverages spatially-regularized non-negative matrix factorization (NMF)^26^ to define individual-specific atlases of functional brain network organization (**Figure 1a**)^3,27^. This approach has been implemented in previous studies using this dataset^18,25^ to identify seventeen PFNs, revealing substantial inter-individual differences in the spatial layout of functional brain regions, with greatest heterogeneity in association networks (**Figure 1b**).

**Figure 1.**
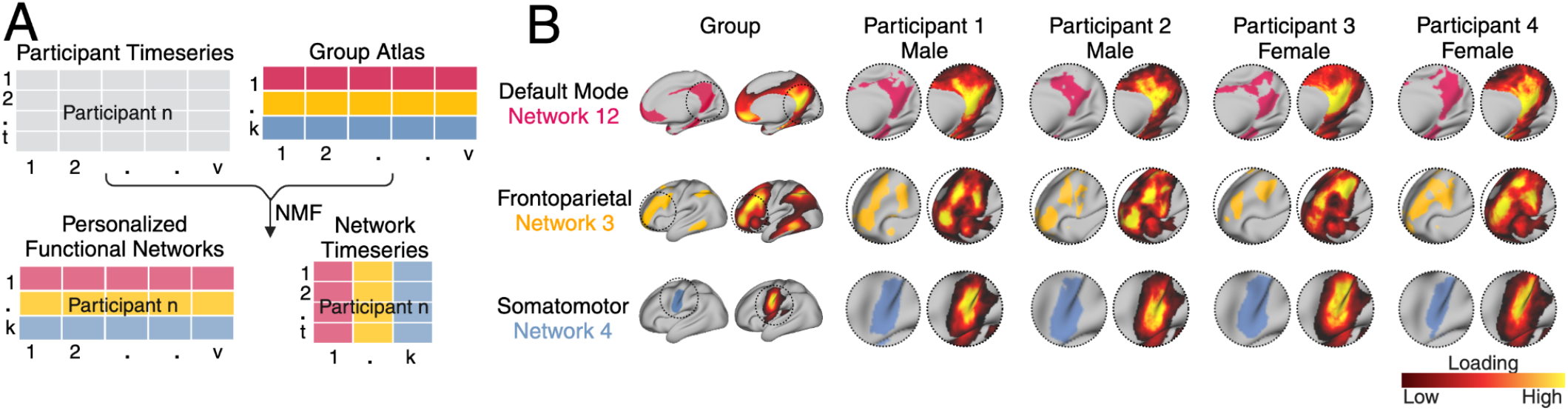
Definition of personalized functional networks (PFNs). *(A)* We employed a precision brain mapping approach that leverages spatially-regularized non-negative matrix factorization (NMF) to define individual-specific atlases of functional brain network organization. In this approach, NMF is performed using a previously-derived group consensus atlas (17 x 59,412) and each individual’s fMRI timeseries. This yields a 17 x 59,412 loading matrix for each participant where each row represents a network, each column represents a vertex, and each cell represents the extent to which each vertex belongs to a given network. This probabilistic definition can be converted into discrete network definitions for display by labeling each vertex according to its highest loading. *(B)* Probabilistic and discrete parcellations of three networks are displayed for the group average and four randomly selected participants. PFNs capture distinct interindividual differences in topographic features. Interindividual variation in topographic features is particularly prominent in association networks such as the default mode network and frontoparietal network. In contrast, sensory and motor networks are more consistent between individuals.

### Mass univariate analysis

To determine whether sex is associated with distinct patterns of PFN topography, we first evaluated vertex-wise associations, as in our previous work^12^ using generalized additive models (GAMs) with penalized splines. These GAMs included a linear covariate for in-scanner head motion (mean fractional displacement), a nonlinear covariate for age, and a random effect covariate for data collection site. Spatial maps of GAM loadings were compared across discovery and replication samples using conservative spin-based permutation testing to account for spatial autocorrelation^28^.

### Multivariate classification

To leverage the high-dimensional data from individual-specific patterns of PFN topography across the whole cortex simultaneously, we next trained a linear support vector machine (SVM) to categorize participant sex based on their multivariate PFN loadings matrix. As in prior work^12^, we applied nested two-fold cross-validation (2F-CV), with the inner loop used to determine the optimal tuning parameter *C* to balance model bias and variance, and the outer loop used to estimate model generalizability to held-out data (**Supplemental Information**). Classifier performance was evaluated using accuracy, sensitivity, specificity, and the area under the receiver operating characteristic (ROC) curve. We also evaluated classifier performance relative to a set of 1000 null models, where participant sex was permuted relative to PFN topography on each iteration, and quantified the relative importance of each feature within the SVM model (**Supplemental Information**).

### Gene expression analysis

After identifying sex differences in PFN topography, we next investigated the biological basis of these differences using imaging transcriptomics. To examine whether sex differences in PFN topography align with cortical gene expression patterns, we compared a summary measure of the overall impact of sex on network loadings across networks—derived from the vertex-wise mass univariate analysis described above—with gene expression data from the Allen Human Brain Atlas^29^. Microarray gene expression data for 12,986 genes were downloaded from https://figshare.com/articles/dataset/AHBAdata/6852911 in the Schaefer1000 atlas parcellation as previously^12^ (**Supplemental Information**).

## Results

We aimed to characterize sex differences in functional brain network topography in a large-scale sample of youth just prior to the transition from childhood to adolescence (*n*=6,437, 9-10 years old, 49.8% female). Given that there are well-documented individual differences in functional topography ^13–15,27^—the spatial layout of functional brain regions across the cortex—we leveraged previously defined maps of personalized functional networks (PFNs; **Figure 1**) for each individual in the ABCD Study^®^ dataset^18^. These maps reflect each individual’s unique functional topography of seventeen canonical large-scale networks.

### Association between sex and person-specific functional topography

To determine whether a participant’s sex is reflected in their person-specific patterns of functional brain network organization, we first conducted mass univariate analyses using generalized additive models (GAMs) to relate vertex-wise PFN loadings to sex. As in our prior work^12^, we fit a GAM at each vertex, including covariates for in-scanner head motion, nonlinear age effects, and site modeled as a random effect, and accounting for multiple comparisons within each PFN by controlling the false discovery rate (FDR; Q<0.05). We found spatially heterogeneous associations between sex and PFN topography in both discovery and replication samples. Sex differences in functional topography were greatest in association networks (**Figure 2A-C; Figure S2-3**), with some PFNs exhibiting greater loadings in females (e.g., fronto-parietal and dorsal attention networks) and others exhibiting greater loadings in males (e.g., default mode and ventral attention networks). We evaluated the total effect of sex at each vertex by summing the absolute value of the Z-statistic across all 17 PFNs. This analysis revealed that associations between sex and PFN topography are greatest in association cortices such as the inferior parietal lobule, ventrolateral prefrontal cortex, and orbitofrontal cortex (**Figure 2D; Figure S4**). We observed highly consistent spatial distributions of GAM loadings across discovery and replication samples (*r*=0.90, *p*_spin_<0.001; **Figure 2E**) and with our prior work in an independent dataset^12^ (*r*=0.59, *p*_spin_=0.0009; **Figure 2F**), using conservative spin-based spatial randomization testing to account for spatial autocorrelation^28^.

**Figure 2.**
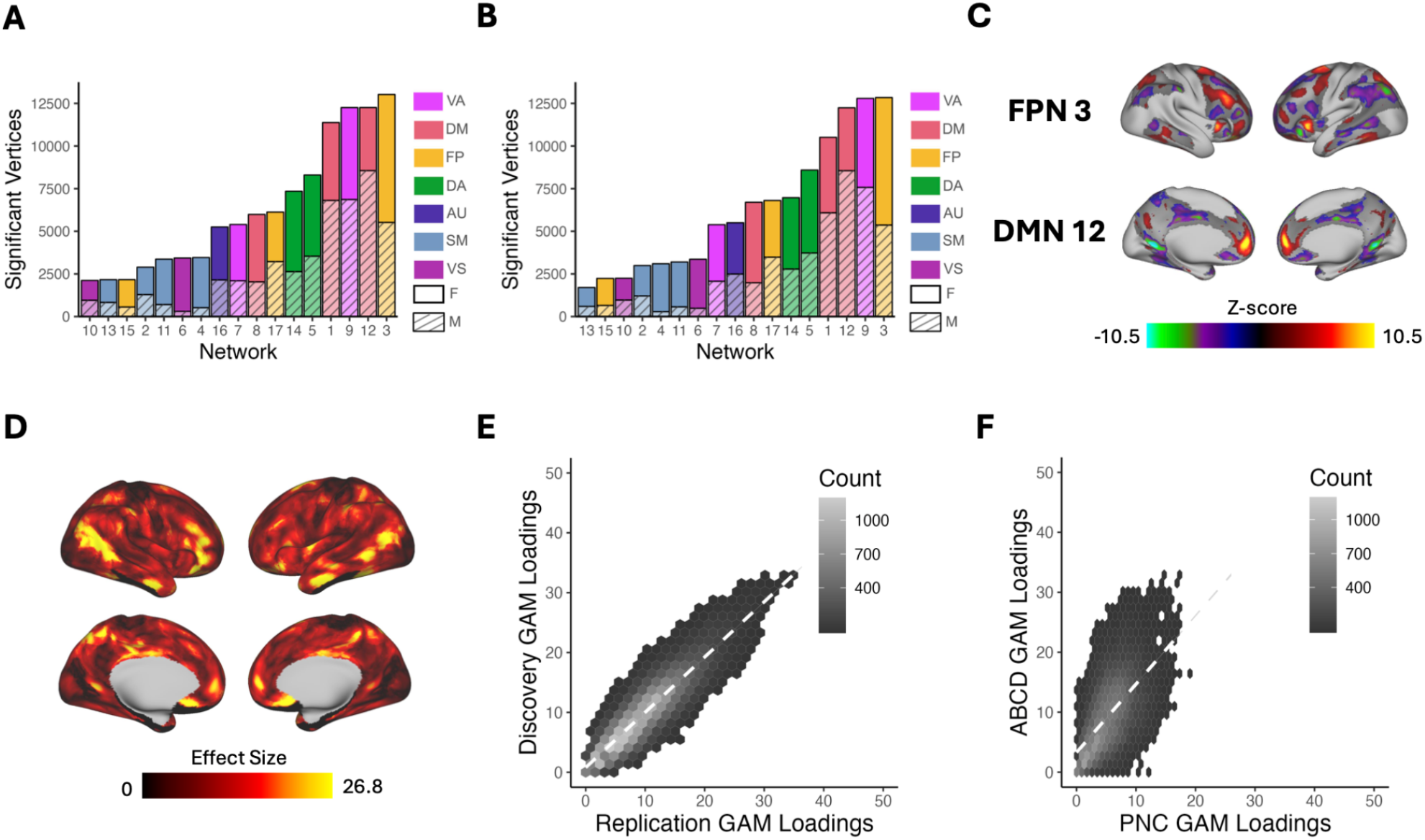
Univariate analysis identifies sex differences greatest in association networks. We fit a generalized additive model (GAM) at each vertex to determine the impact of sex on network loadings. Site, age, and motion were included as covariates with age modeled using a penalized spine and site modeled as a random effect. We accounted for multiple comparisons within each network with false discovery rate (*Q*<0.05). *(A)* The number of vertices in each network with significant sex effects were summed separately for males and females within the discovery set. This process revealed that sex differences were greatest in the association cortex, specifically the frontoparietal, default mode, and the ventral attention networks. *(B)* The same analysis was conducted within the replication set, which yielded convergent results identifying the same 3 networks as having the greatest sex differences. *(C)* Significant vertices are displayed for the frontoparietal, ventral attention, and default mode networks from the discovery set. *(D)* The absolute sex effect across 17 networks was summed to examine the overall effect of sex at a given vertex. The summary measure depicted from the discovery set shows that the areas with the greatest sex effects are in the association cortex. *(E)* The hexplot shows agreement between discovery and replication samples in the association between sex and network loadings (*r* = 0.90, *p*_spin_ < 0.001). *(F)* The hexplot shows agreement between the discovery sample in the ABCD Study and an independent dataset (Philadelphia Neurodevelopmental Cohort; PNC) from our prior report^10^ (*r* = 0.59, *p*_spin_ < 0.0009) in the associations between sex and network loadings. *Abbreviations*: FP/FPN = Fronto-Parietal Network; VA = Ventral Attention; DA = Dorsal Attention; DM/DMN = Default Mode Network; AU = Auditory; SM = Somatomotor; VS = Visual; F = Female; M = Male.

Next, we sought to confirm these vertex-wise mass univariate results by using multivariate classification to leverage the full pattern of PFN topography across the cortex. To evaluate how multidimensional patterns of PFN topography relate to sex, we trained linear support vector machine (SVM) classifiers to categorize participants’ sex from PFN topography patterns using conservative cross-validation. These models were able to correctly identify held-out participants’ sex as male or female from PFN topography patterns with high accuracy averaged across the 100 SVM iterations within each subsample (discovery: 87.4%, replication 87.1%; **Figure 3A, Figure S5A**), successfully replicating our prior work^12^. Model sensitivity and specificity were 0.876 and 0.871, respectively in the discovery sample (replication: 0.869 and 0.869), with a large area under the ROC curve (discovery: 0.966; replication: 0.964), indicating excellent model performance on held-out data that exceeded chance-level accuracy from randomly permuted null models (mean: 0.50, *p*<0.001; **Figure 3A, Figure S5A**, inset histograms).

**Figure 3.**
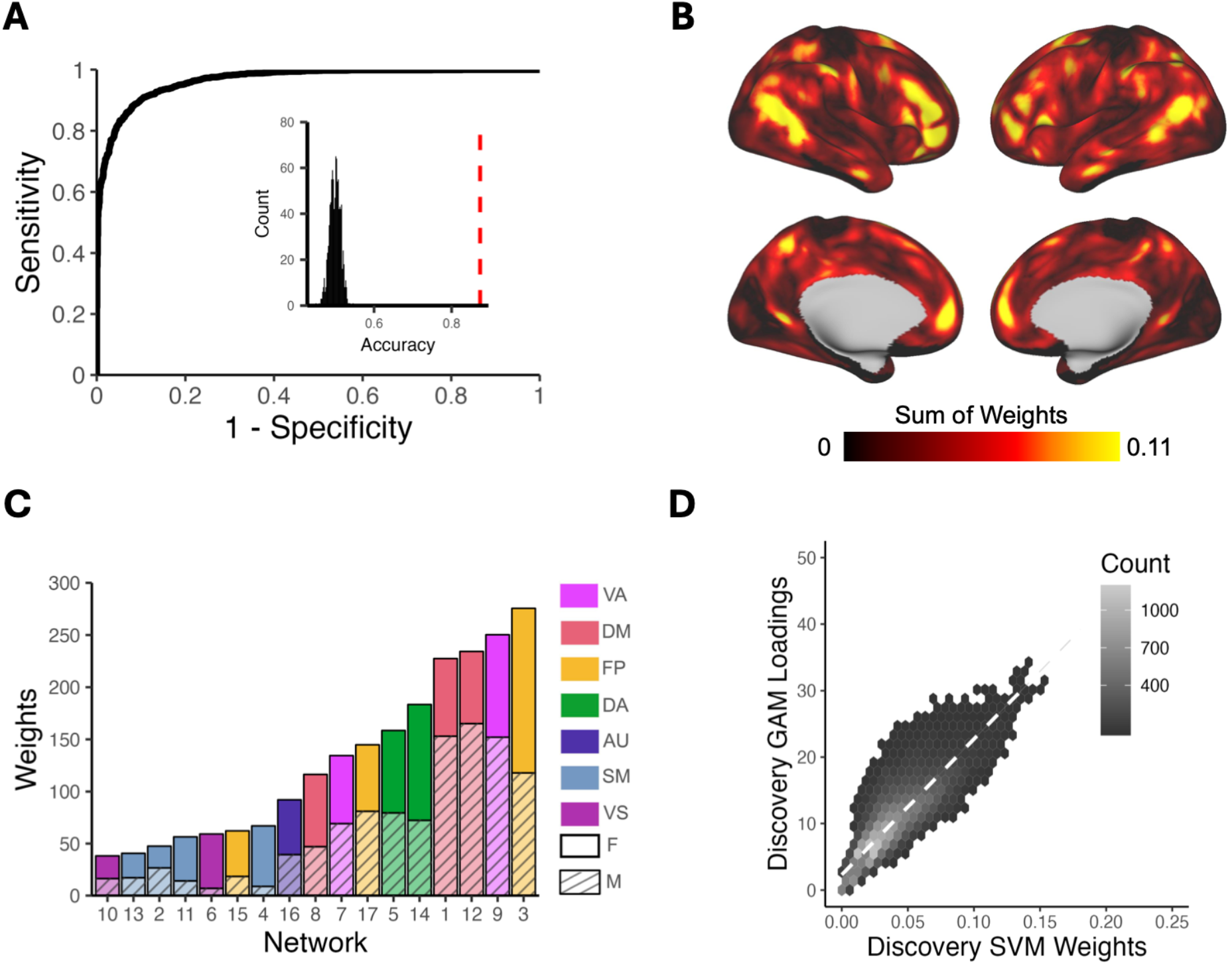
Support vector machine (SVM) models classify participant sex based on PFN functional topography. SVMs were trained with nested two-fold cross-validation (2F-CV) to classify participants’ sex (male or female) from PFN functional topography. *(A)* Depiction of the average ROC curve from 100 SVM models with permuted split-half train-test participant assignments. Average area under the ROC curve was 0.96; average sensitivity and specificity were 0.88 and 0.87, respectively. Inset histogram shows the null distribution of classification accuracies where participant sex was randomized, with the average accuracy from true (non-randomized) data represented by the dashed red line. (*B)* The absolute value of the feature weights were summed at each location across the cortex, revealing that association cortices contributed most to the classification of sex. (*C)* Positive and negative feature weights were summed separately across all vertices in each network to identify which networks contributed most to the classification. Association networks, namely the fronto-parietal, ventral attention, and default mode networks, were identified as the most important contributors for classification. *(D)* Hexplot shows agreement between the absolute summed weights from the multivariate SVM analysis and loadings from the mass univariate generalized additive model (GAM) analysis in the discovery sample (*r* = 0.85; *p*_spin_ < 0.0009). All panels in Figure 3 represent results from the discovery sample. See **Figure S5** for results from the replication sample and **Figure S6** for comparison of SVM weights between the discovery and replication samples. *Abbreviations*: FP = Fronto-Parietal; VA = Ventral Attention; DA = Dorsal Attention; DM = Default Mode; AU = Auditory; SM = Somatomotor; VS = Visual; F = Female; M = Male.

Model performance was robust to the choice of split in participants between the training and testing sets, as evidenced by repeated random cross-validation (discovery: mean accuracy = 87.4% ; 95% CIs = [0.873, 0.874]; replication: mean accuracy = 87.1% ; 95% CIs = [0.870, 0.872]). To identify which brain regions contributed most to the correct classification of participant sex from functional topography, we examined the SVM feature weights after applying the Haufe transformation^30^ to invert the models for interpretability. Replicating prior results^12^, we found that association networks contributed most to the classification of participant sex, primarily those within the fronto-parietal, ventral attention, and default mode networks (**Figure 3B-C; Figure S5B-C**). Vertex-wise patterns of feature weights also provided convergent results with mass univariate analyses (discovery: *r*=0.85, *p*_spin_=0.0009; replication: *r*=0.82, *p*_spin_=0.0009; **Figure 3D; Figure S5D**). The spatial pattern of feature weights was also highly consistent across samples (*r*=0.93, *p*_spin_=0.0009; **Figure S6**).

### Sex differences in personalized functional brain network topography align with X chromosome gene expression patterns

We next sought to investigate genetic correlates of the observed sex differences in person-specific patterns of functional brain network organization. Based on our prior work^12^, we hypothesized that sex differences in PFN topography would align with cortical patterns of sex chromosome gene expression. We therefore conducted a chromosomal enrichment analysis by comparing gene expression data from the Allen Human Brain Atlas (**Supplemental Information**) with the map of vertex-wise sex effects on functional topography from our mass univariate GAM analysis (i.e., the maps in **Figure 2D** and **Figure S4**). We found a significant enrichment of X-chromosome genes in brain regions exhibiting the greatest sex effects in both discovery (*p=*0.004; **Figure 4A**) and replication (*p=*0.003; **Figure S7A**) samples that remained significant when using an FDR-corrected map of vertex-wise sex effects (discovery *p=*0.005; replication *p=*0.004).

**Figure 4.**
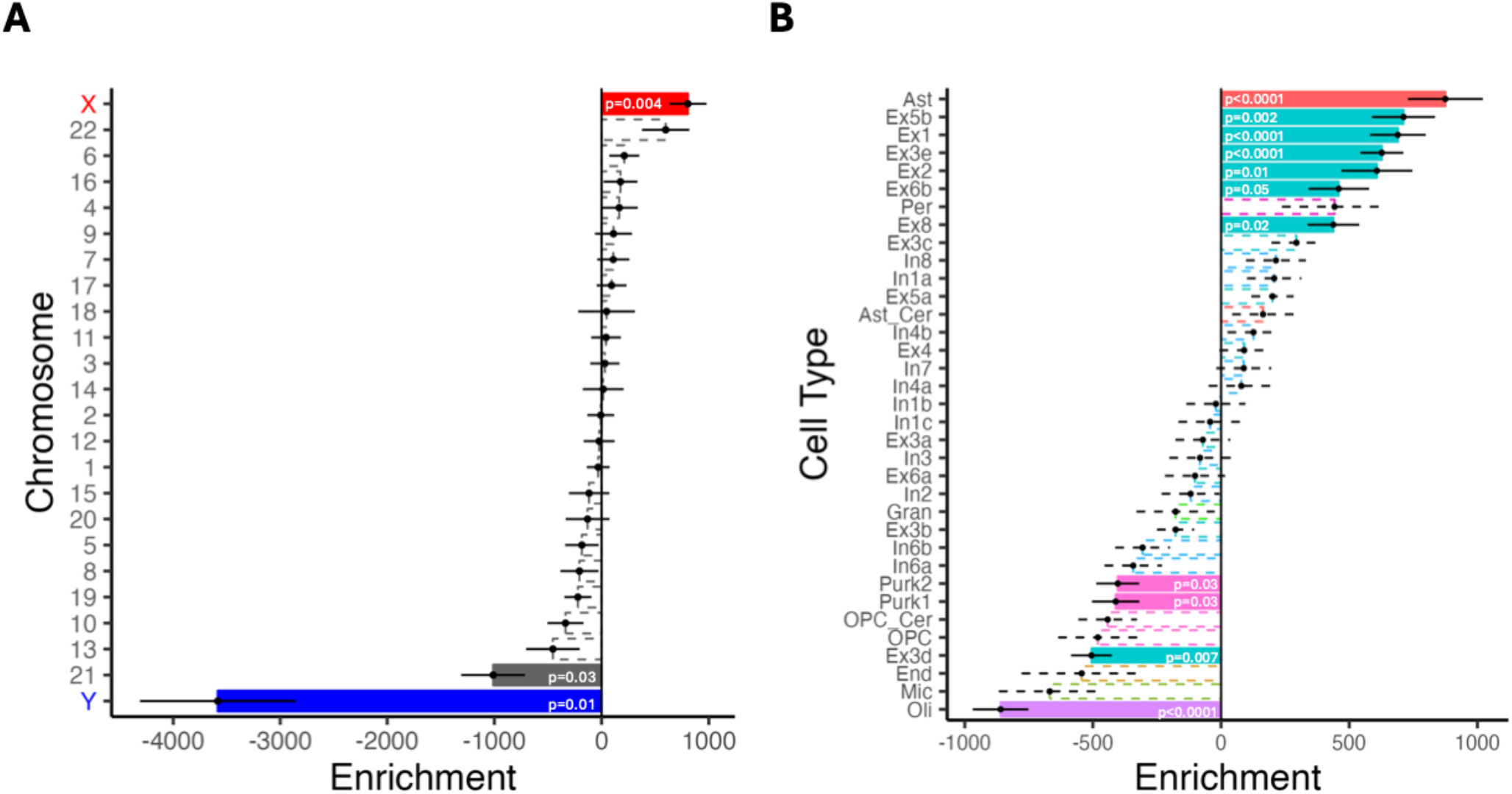
Sex differences in PFN topography align with X-linked and cell-type-specific gene expression patterns. We compared the absolute summed Z-scores from our univariate models to gene expression data from the Allen Human Brain Atlas parcellated to the Schaefer1000 atlas. Point range plots show the median and SE rank of each chromosomal or cell-type gene set. Nonsignificant enrichments are shown by the dashed lines. *(A)* Cortical areas with the greatest sex differences in functional topography were enriched in expression of X-linked genes. *(B)* Cell-type-specific enrichment analyses with cell types assigned via the neuronal subclass assignments determined by Lake et al.^32^ Regions with prominent sex differences in PFN topography were enriched in gene sets related to astrocytes and excitatory neurons, such as Ast, Ex5b, Ex1, Ex3e, and Ex2. *Abbreviations*: Ast = astrocyte; Ast_cer = cerebellar-specific astrocytes; End = endothelial cells; Ex = excitatory neuron; Gran = cerebellar granule cells; In = inhibitory neuron; Mic = microglia; Oli = oligodendrocytes; OPC = oligodendrocyte progenitor cells; OPC_cer = cerebellar-specific oligodendrocyte progenitor cells; Per = pericytes; Purk = cerebellar Purkinje cells.

Among the top ten X-linked genes whose enrichment showed the strongest spatial overlap with sex differences in PFN topography across both the discovery and replication samples, eight genes replicated those found among the top ten in our previous study^12^: *CLCN5, DACH2, GABRA3, GYG2, MUM1L, PCDH19, PDK3, PPEF1*. While it is notable that we observed opposite effects of Y-chromosome genes relative to X-chromosome genes, the scientific significance of the negatively signed Y enrichment effect is difficult to interpret given that the imaging input in our analysis was an absolute value (i.e., an absolute value of the effect of sex summed across networks).

Given that regional differences in cellular composition may be reflected in the spatial patterning of cortical gene expression^31^, we next conducted cell-type-specific gene expression analysis. First, using cell-type specific gene sets as assigned by Seidlitz et al.^31^, we found that regions exhibiting the greatest sex differences in PFN topography were enriched in gene expression of astrocytic (discovery: *p*<0.0001; replication: *p*<0.0001) and excitatory neuron (discovery: *p*<0.0001; replication: *p*<0.0001) cell types. These enrichments remained significant when using an FDR-corrected map of vertex-wise sex effects in both the discovery and replication samples (discovery: *p*<0.0001; replication: *p*<0.0001). We found convergent results using a finer-grained cell type subclass assignments^32^ (**Figure 4B; Figure S7B**): regions showing the greatest sex differences in PFN topography exhibited enrichment for astrocytic gene expression (*p*<0.0001) and several excitatory neuron subtypes such as Ex5b (*p*=0.002), Ex1 (*p*<0.0001), Ex3e (*p*<0.0001), Ex2 (*p*=0.014), Ex6b (*p*=0.047), and Ex8 (*p*=0.021), demonstrating a strong replication of our prior work in an independent dataset^12^.

## Discussion

Our results demonstrate robust and replicable associations between sex and the spatial patterning of functional brain networks in youth. Across analytic approaches and independent samples, we consistently find that the spatial patterning of person-specific functional brain networks significantly differs based on sex as a biological variable. While no single brain region or network is systematically larger or smaller in its spatial extent across all males or females, we find that the greatest sex differences in functional topography tend to be disproportionately found in association areas like the fronto-parietal, default mode, and ventral attention networks, with weaker effects found in sensory and motor cortices. We also found that the spatial distribution of sex differences across the cortex aligns with X-linked gene expression patterns, as well as signatures of astrocytic and excitatory neuronal cell types. These results suggest that sex might be one of many factors that shape the development of functional networks in youth.

### Sex differences in personalized functional brain network topography in youth

Extending prior work in this area, our results suggest that sex differences in functional topography are consistently observed in children just prior to the transition to adolescence. This critical transition period that often coincides with pubertal changes is marked by the emergence of many common psychiatric disorders, including depression and anxiety, which disproportionately affect females^1^. This time period also coincides with the maturation of functional brain networks, including the protracted topographical refinement of association networks like the fronto-parietal and default mode networks^33^. These association networks also exhibit the most person-specific patterns of functional topography among all large-scale brain networks and are associated with symptoms of psychopathology^16^. Our observation that these networks also reflect an individual’s sex aligns with recent findings^34^ and suggests that sex differences in functional brain networks may play a role in the emergence and exacerbation of sex differences in psychiatric disorders during the transition to adolescence. Thus, future studies may seek to further investigate the potential role of functional brain network development as an early biomarker for sex-specific psychiatric symptom emergence in youth.

### Sex differences in PFN topography are spatially correlated with X-linked gene expression

The present results highlight potential biological mechanisms by which sex might shape the spatial patterning of functional brain networks. By comparing patterns of sex-linked gene expression with the spatial distribution of sex differences in PFN topography, we found consistent overlap with X-chromosome gene expression, suggesting that sex differences in functional brain network architecture may, in part, result from differential gene expression patterns. Strikingly, among the 474 possible X-linked genes included in processed Allen Human Brain Atlas microarray data^35^, we found that the same short list of eight genes was consistently ranked in the top ten for the highest spatial correspondence with sex differences in functional topography across the discovery sample, the replication sample, and an independent study^12^.

Several of these genes have been suggested to be important for neuronal structure, function, metabolism, and development^36–38^. For example, one highly-ranked X-linked gene in particular, *PCDH19*, encodes a protocadherin protein that plays a critical role in neurodevelopment by supporting neuronal organization and migration. Moreover, mutations in this gene are associated with neurodevelopmental disorders such as autism spectrum disorder^37^. While multiple mechanisms are likely at play, our observation that *PCDH19* is consistently associated with sex differences in person-specific functional brain network organization across diverse samples of youth highlights the potential importance of further investigating this pathway in sex-specific neurodevelopmental trajectories and psychiatric illness.

As in prior work, the present results suggest that functional brain network configuration may also be shaped by the organizational effects of sex hormones on cytoarchitecture. In line with our previous findings, our results show that cortical areas with the greatest sex differences align with genes expressed in excitatory neurons and astrocytes. This result is particularly salient in the context of the vast nonhuman primate and rodent literature demonstrating the effects of estradiol on glutamatergic dendritic spine architecture^39,40^ as well as astrocytic structure, proliferation, and apoptosis^41,42^. These results have now been replicated using two different cell-type categorizations in three distinct samples across two independent datasets, motivating future research into cytoarchitectural mechanisms.

### Sex differences in functional topography and alignment with gene expression patterns consistently replicate across independent datasets

Replication studies often fail^43^, and even successful replication studies most often yield results with smaller effect sizes than initial discoveries^44^. The present study not only successfully replicates *all* findings tested in our prior work, it also uncovered effect sizes that were approximately the same or even larger than in the previous study^12^. Specifically, the present study confirmed the presence of sex differences in PFN topography and replicated the observation that these sex differences are primarily found in association networks. We also found a striking replication in our imaging transcriptomic analyses, with spatial patterns of gene expression in the same astrocytic and neuronal cell types showing overlap with brain regions exhibiting the greatest sex differences. This definitive replication is especially notable in light of the many differences between the datasets in each study, including sample size, age range, scanner types and protocols, data collection sites, functional MRI tasks, racial/ethnic diversity, and socioeconomic status. Thus, the present study represents a strong counterexample to the ongoing reproducibility crisis in psychology and neuroscience^19^.

Several important distinctions between the present study and this previous work provide context for interpreting these results. First, the previous study^12^ used data from the Philadelphia Neurodevelopmental Cohort (PNC; *n*=693). Here, we applied the same analytical approach to a dataset that is an order of magnitude larger (ABCD Study^®^; *n*=6,437). This considerable increase in sample size may explain the improvement in model performance on held-out data between studies (from 82.9% to 87.1% accuracy), as models trained in larger datasets with rigorous cross-validation are less likely to be overfit^45,46^. Second, the previous study^12^ assessed individuals aged 8-23 years old, while the present study leveraged data from the baseline assessment of the ABCD Study^®^ when participants were 9-10 years old. The more restricted age range in the present study may also help to explain the improved model performance, since functional brain network topography changes throughout development^2,3^. Though age was included as a model covariate in both studies, it is possible that the smaller age range in the present study still yielded some advantage in classifying sex from patterns of functional topography at a more restricted time period of brain development.

### Limitations

There are several limitations of this study worth noting. First, sex was assessed using a binary parent-reported question regarding the assignment of sex at birth on the original birth certificate, and we lacked a sufficiently large sample size to examine functional topography of intersex youth. Importantly, existing data suggest that binary classifications of sex do not align well with the complex mosaics of male and female characteristics observed in individual brains^47^. Thus, further research is warranted to more comprehensively characterize person-specific patterns of male, female, and intersex characteristics in functional brain network topography. Second, prior work has shown that functional brain network connectivity is associated with both sex and gender in youth^48^. As the present study aimed to understand sex differences in functional topography, future work is also needed to investigate potential effects of continuous gender dimensions such as gender identity and expression. Given that only 0.5% (*n*=58) of baseline ABCD Study^®^ participants reported being or possibly being transgender^49^, and given that gender continues to develop throughout early adolescence, future studies in longitudinal timepoints will be key in investigating potential individual or interactive effects of sex and gender in shaping neurodevelopment.

Third, the present study leveraged a cross-sectional sample at a single time point from within an ongoing longitudinal study of youth. As youth from the ABCD Study^®^ continue to participate in follow-up study sessions from childhood to adulthood, it will become increasingly possible to link changes in sex-specific functional brain network topography with critical developmental changes such as puberty. Future longitudinal studies considering the complex interplay of biopsychosocial factors related to sex and gender development may also reveal mechanistic links between sex-specific patterns of functional brain network topography and sex differences in psychiatric illness manifestation (e.g., internalizing symptoms). Fourth, the present study focused on sex differences in functional rather than structural differences in brain organization, though sex differences in gross structural anatomy (e.g. head size) are well documented^50^. However, recent work has demonstrated that sex differences in functional brain organization do not appear to be systematically associated with structural imaging measures such as surface area or microstructural organization^34^. Finally, as previously noted^12^, there are several limitations associated with use of the Allen Human Brain Atlas, including donor age and sex, gene expression quantification, asymmetric sampling, and sample size. Importantly for the present study, the Allen Human Brain Atlas includes postmortem samples from five male donors and one female donor, so replication of these findings in a sex-balanced sample is warranted when such spatially comprehensive maps of gene expression become available.

### Future directions: using precision brain mapping to inform female mental health

In addition to the future directions noted above, our observation that person-specific patterns of functional brain network topography show sex differences, particularly in association networks related to psychiatric symptoms^17^, also lays important groundwork for future studies of female mental health. First, future work should further examine how PFN topography develops across the female reproductive lifespan, with a particular focus on changes across critical hormonal transition periods such as puberty, pregnancy, and menopause. These hormonal transition periods are known to have substantial impact on neurodevelopment and often align with the timing of psychiatric illness onset^51^, yet have been historically underfunded and understudied^52^. Extending the study of PFNs across the lifespan therefore has potential to improve our understanding of how neuroplasticity during hormonal shifts impacts functional topography and trajectories of psychiatric illness.

Second, longitudinal studies examining how sex differences in PFN topography emerge during development may inform early preventions or personalized treatments such as neuromodulation via transcranial magnetic stimulation (TMS), filling critical gaps in existing treatment options. Finally, it is worth noting that these sex-specific individual differences in the topography of personalized functional brain networks are also strongly associated with childhood environments and socio-economic status^25^. These factors have been shown to confer vulnerability to psychiatric symptoms during future reproductive timepoints characterized by significant hormonal fluctuations such as pregnancy^53^ and menopause^54,55^. Future work may therefore seek to parse the effects of hormonal, genetic, and environmental factors that may together shape individual-specific spatial patterning of functional networks across the female reproductive lifespan.

## Conclusions

In this study, we demonstrate reproducible sex differences in person-specific patterns of functional brain network organization in youth. The ability to successfully classify sex from the spatial configuration of PFNs is primarily driven by sex differences in the functional topography of association networks. Brain areas that are most strongly associated with sex are also enriched in the expression of X-linked genes and genes expressed in excitatory neurons and astrocytes, highlighting a potential genetic basis for sex differences in functional brain network topography. By characterizing sex differences in functional topography in youth, this study provides a key stepping stone toward addressing sex differences in susceptibility to psychiatric symptoms that emerge during the transition to adolescence.

## Data Availability Statement

Data used in the preparation of this article were obtained from the Adolescent Brain Cognitive Development Study® (https://abcdstudy.org), held in the NIMH Data Archive (NDA). Only researchers with an approved NDA Data Use Certification (DUC) may obtain ABCD Study data.

## Analytic Code Availability Statement

Analytic code used in this study is available at https://ashleychari.github.io/abcd_sex_pfn_replication/

## Author Contributions

SS, TDS, AAB, and JS formulated the research questions. ASK, ZC, JS, AAB, TDS, and SS designed the study. ASK, KYS, AF, ZC, ARP, and SS performed the research including analyzing the data. ASK, AF, HR, EB, and SS drafted the first draft of the work. ASK, KYS, AF, HR, EB, DSB, MC, ZC, CD, YF, MG, RK, SK, BL, HL, IL, ARP, LP, AR, DRR, JS, GS, RTS, DHW, AAB, TDS, and SS interpreted data, and reviewed and revised the work critically for intellectual content.

