## Supplemental Information for "Reproducible Sex Differences in Personalized Functional Network Topography in Youth"

### Supplementary Information

#### Supplementary Methods

**Functional Magnetic Resonance Imaging (fMRI) Processing:** As in our prior work[15,20], we leveraged data from the ABCC Collection 3165 processed with the ABCD-BIDS pipeline[17] which included distortion correction and alignment, Advanced Normalization Tools (ANTS[21]) denoising, FreeSurfer[22] segmentation, and surface and volume registration with rigid-body transformation[23]. Following this, further processing was done using the DCAN BOLD Processing (DBP) pipeline which includes de-meaning and de-trending of fMRI data with respect to time, denoising using a general linear model with regressors for signal and movement, bandpass filtering between 0.008 and 0.09 Hz using a 2nd order Butterworth filter, applying the DBP respiratory motion filter (18.582–25.726 breaths per minute), and applying DBP motion censoring (frames exceeding an FD threshold of 0.2 mm or failing to pass outlier detection at +/− 3 standard deviations were discarded). We then concatenated cleaned time series data for resting-state and task-based scans as in previous work[15,20] to maximize the data available for analysis. We excluded participants who had fewer than 600 remaining TRs after motion censoring, as well as those who failed ABCD quality control for their T1 or resting-state fMRI scan.

**Multivariate classification model training and testing:** Prior to model training and testing, we eliminated siblings from training and testing subsamples to avoid leakage of family structure across subsamples, yielding a total sample of $n$ = 6,437 (discovery: $n$ = 3240, 50.46% female;

replication: $n$ = 3197, 49.13% female) for all multivariate classification analyses. Before beginning our 2F-CV procedure, we first split the data between the matched discovery and replication samples according to the previously defined ABCD Reproducible Matched Samples[20,22]. Then, separately within the discovery and replication samples, we performed 2F-CV as follows (see Figure S1 in Shanmugan et al.[12] for a visual depiction of our nested 2F-CV procedure, here applied separately to matched discovery and replication samples). For the outer 2F-CV loop, we trained and tested the SVM model using split-half subsets separately within either the discovery or replication sample. After training the model in one half of the data and testing its performance in the other held-out half of the data, we then repeated this procedure in reverse. Prior to model training, covariates for age, site, and in-scanner head motion (mean fractional displacement) were regressed from each feature, separately in the training and testing sets to avoid leakage. To determine whether classification accuracy was driven by the choice of split, we repeated this analysis using 100 permuted splits of the data, each time randomly dividing the discovery and replication samples into independent training and testing sets.

Inner 2F-CV loops were used to determine the optimal tuning parameter C by further randomly dividing the training set of the outer 2F-CV loop into two subsamples. The first split-half subsample was used to train the SVM model with each of 15 possible C parameter values: [$2^{-5}$, $2^{-4}$, … , $2^{8}$, $2^{9}$]. These models were each tested in the second held-out subsample as in our previous work[12]. We then repeated this procedure using the second subsample for training and the first subsample for testing, calculating the average held-out classification accuracy across the two subsamples for each value of the parameter C. The optimal C parameter value was selected as the C with the highest average held-out classification accuracy, and this optimal C parameter was used to train the models within the outer 2F-CV loop. It is worth noting that even the smallest subdivisions of the data in our nested 2F-CV procedure still contained over one thousand participants each at a minimum, yielding sufficient statistical power to train and test our machine learning models using the most conservative possible (fewest folds) cross-validation approach.

**Evaluation of Feature Importance:** To evaluate the relative importance of each feature within the SVM model, we first extracted feature weights for each network loading at each vertex and averaged these weights across the 100 randomly permuted splits of the data. Then, to avoid challenges with interpretation due to the covariance structure among feature weights, we applied the Haufe transformation[28] to invert the models prior to feature weight interpretation. Next, we averaged the Haufe-transformed weight maps across the training and testing sets from the outer loop of the matched samples 2F-CV procedure. As in our univariate analysis, spatial maps of SVM weights were compared across samples using spin-based permutation testing[27].

**Analysis of gene expression:** Details on gene processing, including sample assignment, probe selection, gene information reannotation, data filtering, data normalization, and gene filtering are described in Arnatkeviciute et al.[30]. Given that just two out of six donor brains contained samples from both the left and right hemispheres, we restricted gene expression and chromosomal enrichment analyses to the left hemisphere, thus ensuring maximum sampling of probes[30]. Consistent with our prior work[12], genes were assigned to each chromosome according to annotation by Richiardi et al.[34], and we calculated the median ranks for 24 non-overlapping gene sets which included autosomal chromosomes 1-22 and sex chromosomes X and Y. We then compared the calculated median rank within each chromosomal gene set to a same-sized null distribution of median ranks calculated from 1,000 non-parametric permutations in which the original ranked list was randomly reordered. The corresponding p-value from this permutation test was calculated as the proportion of permutations with a more extreme value than the median rank of the true data and was not further corrected for multiple comparisons in line with prior studies using these methods[12,31].

As regional differences in the spatial pattern of gene expression may reflect regional differences in the cellular composition of each cortical area[31], we also investigated cell-type-specific gene enrichments to probe the convergent and divergent patterns of discrete underlying gene sets, as in our prior work[12]. We again used ranked gene lists and nonparametric permutation testing as in our chromosomal enrichment analyses to test whether the spatial pattern of a cell-type-specific gene set was non-randomly associated with the spatial pattern of sex differences in functional topography. Gene sets for each cell type were first categorized according to prior work[31]. We then applied previously-determined, finer-grained neuronal subclass assignments[35] to obtain a more nuanced understanding of cytoarchitecture. In both analyses, only brain-expressed genes[31] defined by expression levels in the Human Protein Atlas[36,37] were considered. To test the relationship between the spatial pattern of sex differences in PFN topography and the spatial patterns of gene expression for a given gene set in each ranked gene list, we quantified the degree of spatial correspondence using the median gene set rank as in prior studies[12,31–33].

#### Supplementary Figures

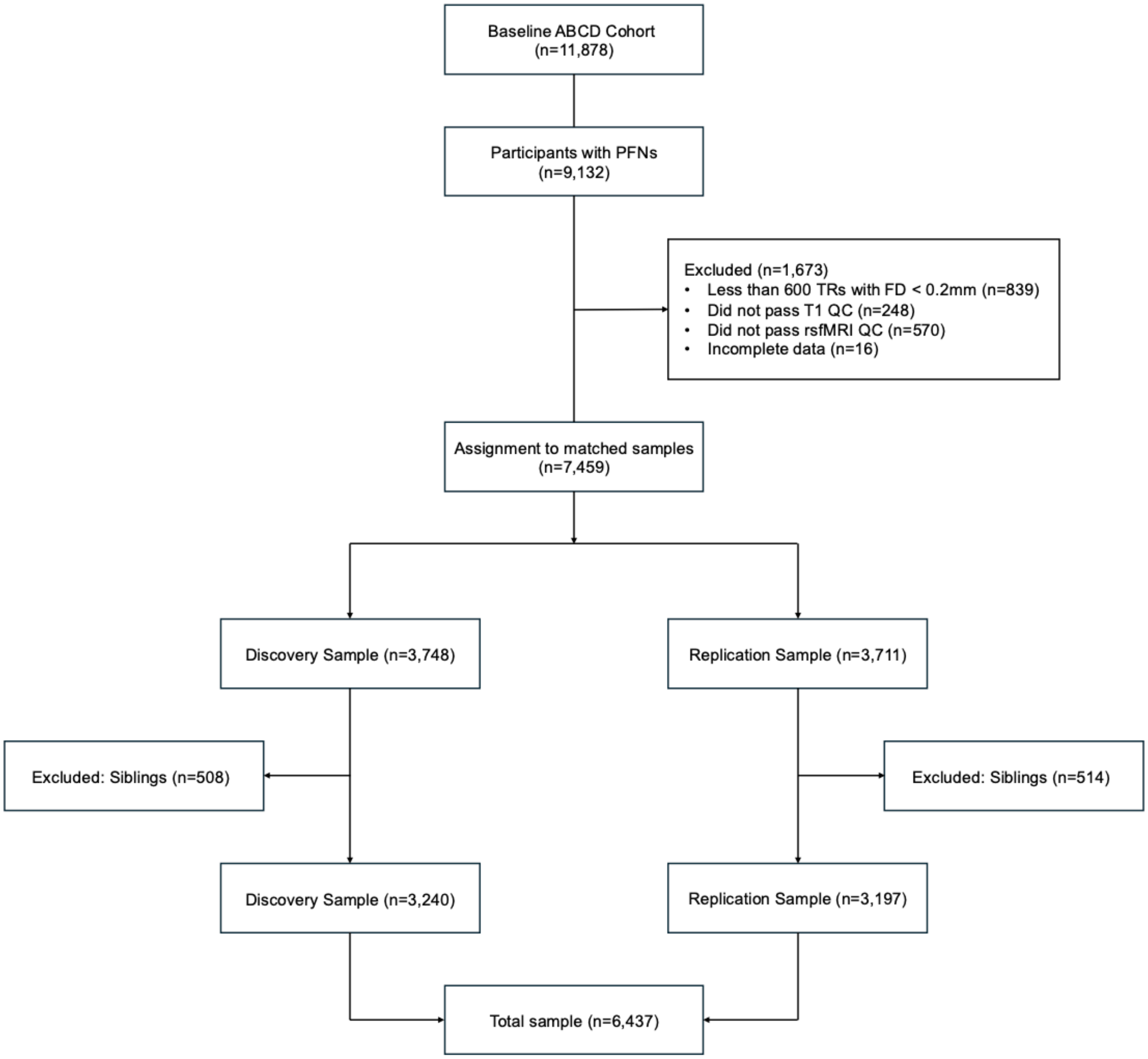


**Figure S1. Flow diagram depicting data inclusion and exclusion.** Participants from the Adolescent Brain Cognitive Development (ABCD) Study®[16] baseline assessment (*n* = 11,878) were drawn from the ABCD BIDS Community Collection (ABCC, ABCD-3165[17]). Participants were excluded for having incomplete data or excessive head motion. The matched samples were then split into discovery and replication samples according to the ABCD Reproducible Matched Samples (ARMS[17]). Siblings were excluded from the discovery and replication sets separately to avoid leakage across subsamples during two-fold cross-validation, yielding a total of *n* = 3,240 participants in the discovery sample and *n* = 3,197 participants in the replication sample.

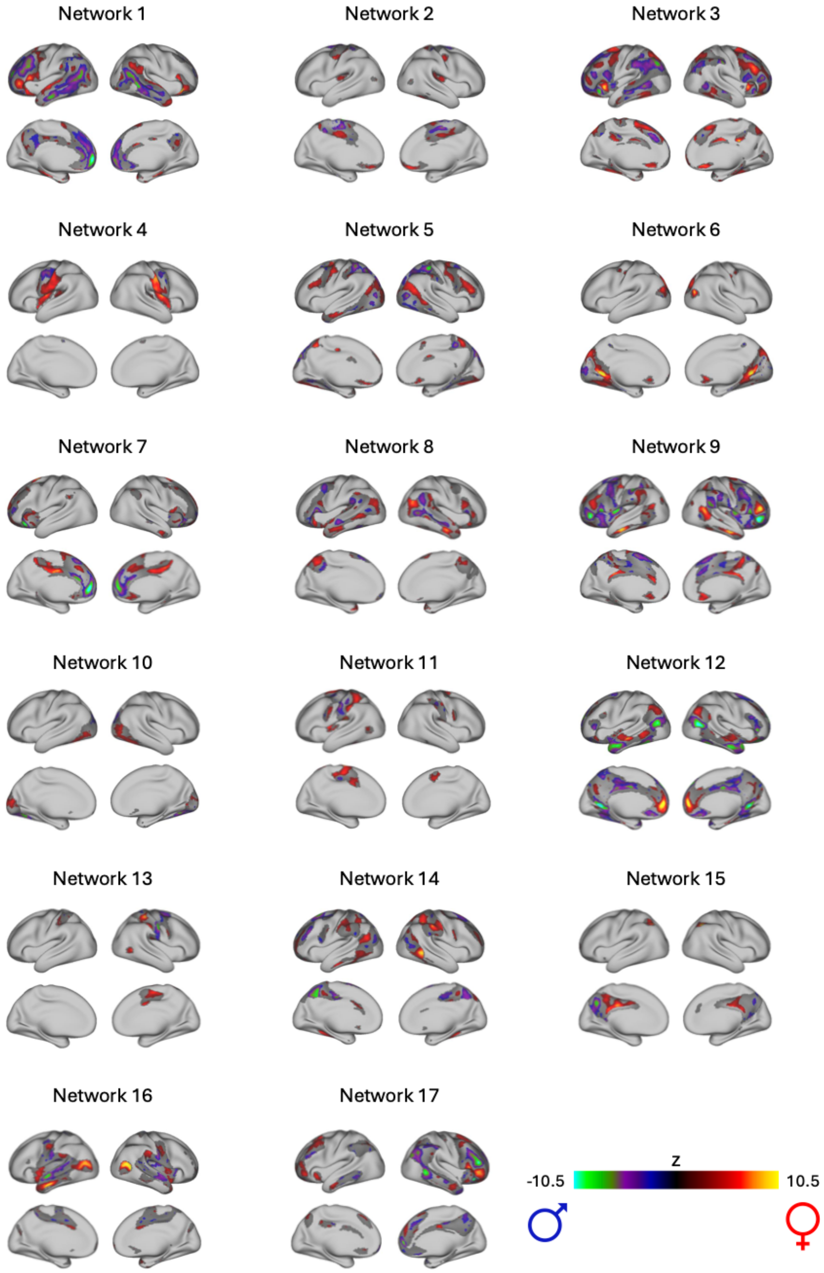


**Figure S2. Univariate analyses identify significant sex differences in association networks.** A GAM was fit at each vertex to evaluate the association between sex and network loadings. Age, site, and head motion were included as covariates with age modeled using a penalized spine and site modeled as a random effect. Multiple comparisons were accounted for by controlling the false discovery rate ($Q$<0.05). Significant vertices are shown for each of the 17 PFNs.

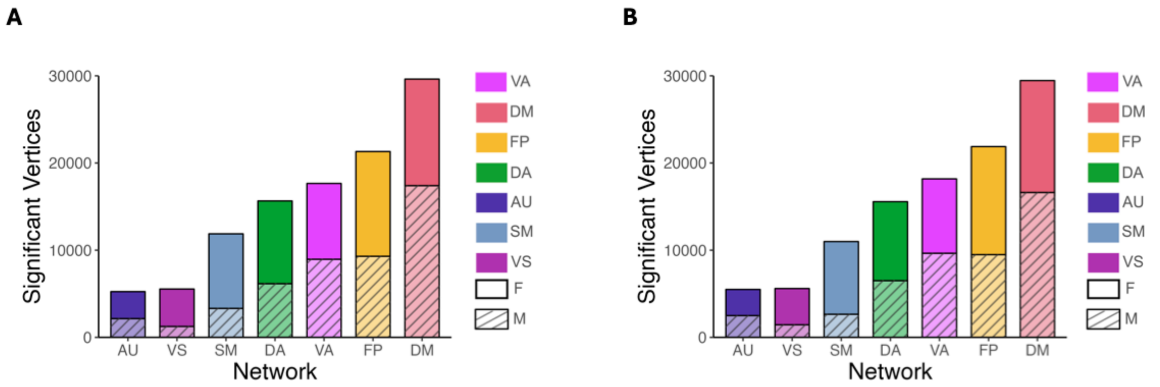


**Figure S3. Univariate analysis identifies significant sex differences in association networks in both discovery dataset and replication dataset.** A GAM was fit at each vertex to evaluate associations between sex and PFN network loadings. Age, site, and motion were included as covariates with age modeled using a penalized spine and site modeled as a random effect. Multiple comparisons within each network were accounted for by controlling the false discovery rate ($Q$<0.05). The number of significant vertices for each network category was summed separately for males and females (e.g., the "DM" bar represents the number of vertices with significant sex effects in networks 1, 8, and 12). Sex differences were greatest in association networks in both the discovery *(A)* and replication *(B)* samples. *Abbreviations*: FP = Fronto-Parietal; VA = Ventral Attention; DA = Dorsal Attention; DM = Default Mode; AU = Auditory; SM = Somatomotor; VS = Visual; F = Female; M = Male.

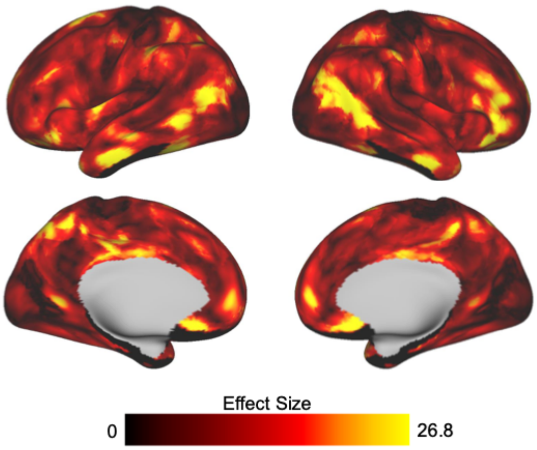


**Figure S4. Univariate analysis identifies significant sex differences in association networks in the replication sample.** We summed the absolute sex effect across 17 networks to examine the overall effect of sex at a given vertex within the replication sample. Brain areas with the greatest sex effects are found in association cortices.

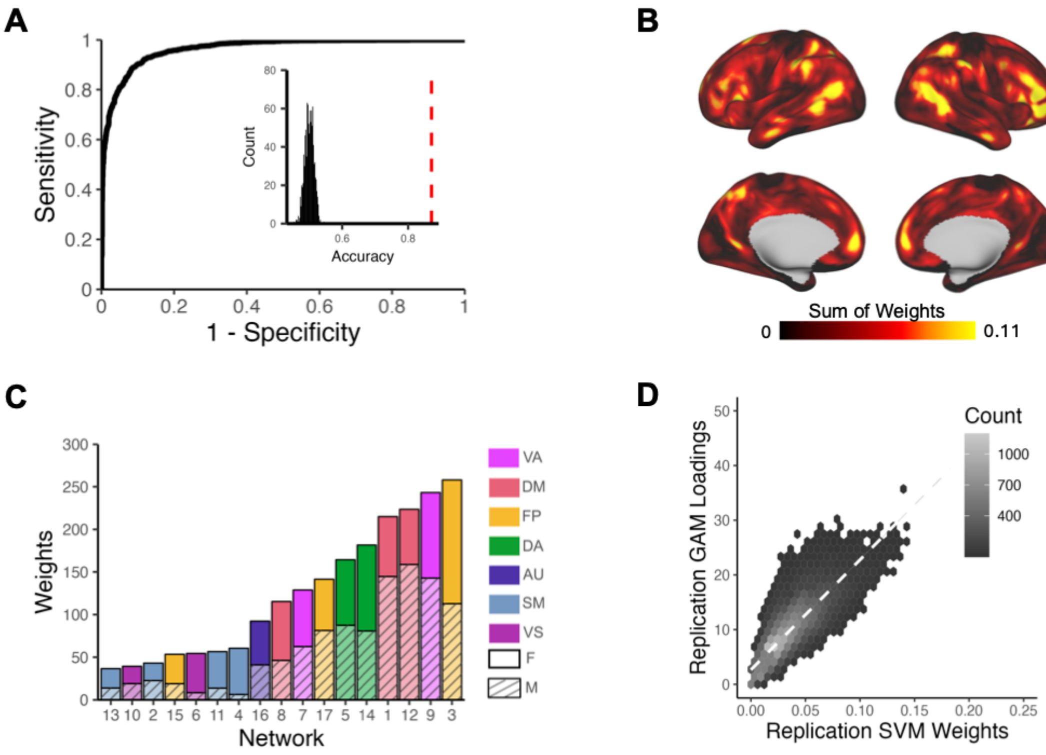


**Figure S5. Support vector machines (SVMs) accurately classify participant sex from PFN functional topography in the replication dataset.** SVMs were trained with nested two-fold cross-validation (2F-CV) to classify participants' sex (male or female) based on PFN functional topography. *(A)* Depiction of the average ROC curve from 100 SVM models with permuted split-half train-test participant assignments. Average area under the ROC curve was 0.96; average sensitivity and specificity were 0.87 and 0.87, respectively. Inset histogram shows the null distribution of classification accuracies where participant sex was randomized, with the average accuracy from true (non-randomized) data represented by the dashed red line. *(B)* The absolute value of the feature weights were summed at each location across the cortex, revealing that association cortices contributed most to the classification of sex. (*C)* Positive and negative feature weights were summed separately across all vertices in each network to identify which networks contributed most to the classification. Association networks, namely the fronto-parietal, ventral attention, and default mode networks, were identified as the most important contributors for classification. *(D)* Hexplot shows agreement between the absolute summed weights from the multivariate SVM analysis and loadings from the mass univariate generalized additive model analysis in the discovery sample ($r = 0.82$; $p_{spin} < 0.001$). *Abbreviations*: FP = Fronto-Parietal; VA = Ventral Attention; DA = Dorsal Attention; DM = Default Mode; AU = Auditory; SM = Somatomotor; VS = Visual; F = Female; M = Male.

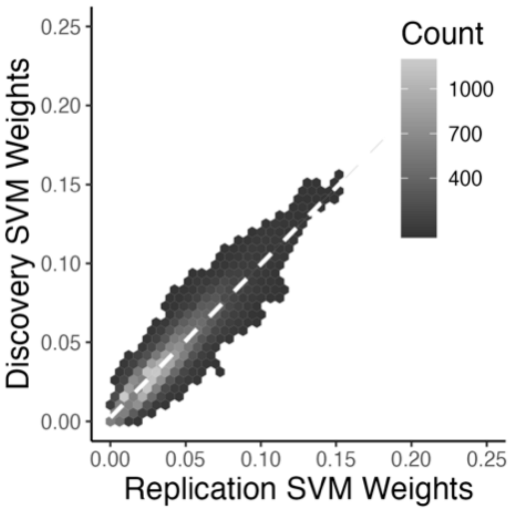


**Figure S6. Comparison of support vector machine (SVM) feature weights across samples.** The hexplot shows agreement between discovery and replication samples in the association between sex and network loadings from the SVM models ($r$ = 0.93; $p_{spin}$ < 0.001).

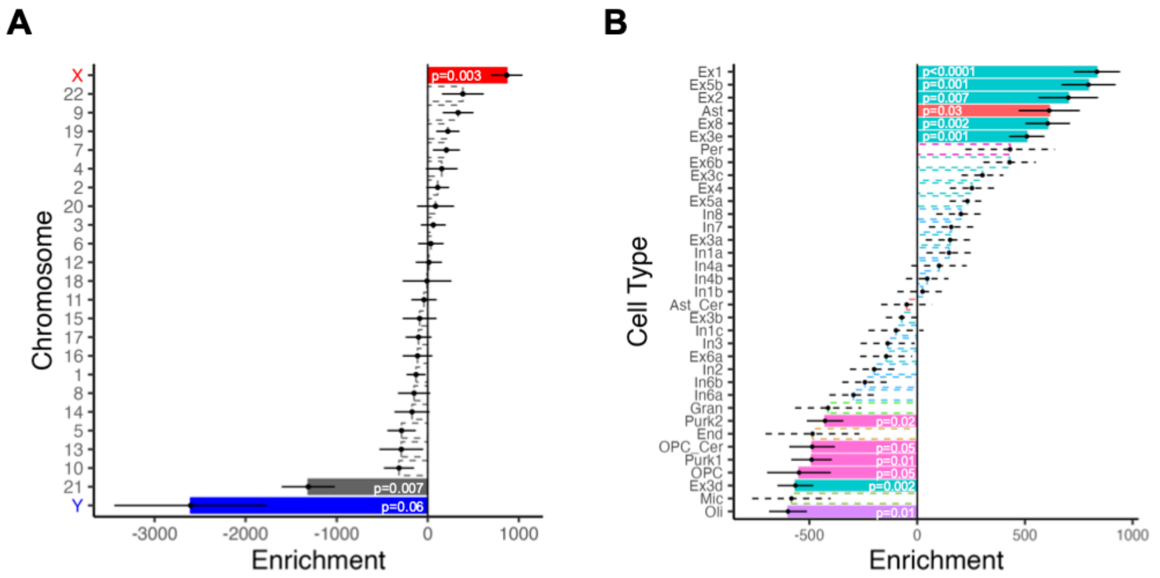


**Figure S7. Alignment between sex differences in PFN topography and expression of X-linked genes within replication dataset.** We compared the absolute summed Z-scores from our univariate models to gene expression data from the Allen Human Brain Atlas parcellated to the Schaefer1000 atlas. Point range plots show the median and SE rank of each chromosomal or cell-type gene set. Nonsignificant enrichments are shown by the dashed lines. *(A)* Cortical areas with the greatest sex differences in functional topography were enriched in expression of X-linked genes. *(B)* Cell-type-specific enrichment analyses with cell types assigned via the neuronal subclass assignments determined by Lake et al.[32] Regions with prominent sex differences in PFN topography were enriched in gene sets related to astrocytes and excitatory neurons, such as Ex1, Ex5b, Ex2, Ast, Ex8, and Ex3e. *Abbreviations*: Ast = astrocyte; Ast_cer = cerebellar-specific astrocytes; End = endothelial cells; Ex = excitatory neuron; Gran = cerebellar granule cells; In = inhibitory neuron; Mic = microglia; Oli = oligodendrocytes; OPC = oligodendrocyte progenitor cells; OPC_Cer = cerebellar-specific oligodendrocyte progenitor cells; Per = pericytes; Purk = cerebellar Purkinje cells.
